# Transient AMPK activation by nutrient stress of high fat diet preserves cardiac electrophysiological stability and protects against arrhythmias

**DOI:** 10.1101/2025.06.11.658631

**Authors:** Michael W. Rudokas, Margaret McKay, Xiaohong Wu, Jonathan Granger, Yancey Williams, Markus Bögner, Taekyung Kang, Anton Seyfried, Zeren Toksoy, Marine Cacheux, Lawrence H. Young, Fadi G. Akar

## Abstract

Sudden cardiac death (SCD) is a major complication of obesity, yet it remains unclear whether early metabolic stress, prior to the onset of overt obesity or structural remodeling, can independently promote arrhythmias. In vitro studies suggest that fatty acids can allosterically stimulate AMP-activated protein kinase (AMPK), a key metabolic sensor known to preserve myocardial viability and mitochondrial function following ischemia-reperfusion (I/R) injury. We hypothesized that AMPK signaling critically modulates the electrophysiological (EP) response to high-fat diet (HFD)-induced metabolic stress.

**Methods:** To test this, wild-type (WT) and AMPK kinase-dead (AMPK-KD) mice were subjected to an 8-week HFD regimen beginning at 4 weeks of age. Controls remained on normal diet (ND) for the same duration. Arrhythmia susceptibility was assessed ex vivo using rapid pacing and I/R challenge protocols. Changes in the EP substrate were defined by high-resolution optical action potential mapping. Underlying mechanisms were probed using western blotting, confocal and transmission electron microscopy.

**Results:** HFD-fed wild-type (WT) hearts did not display increased arrhythmia susceptibility in response to either burst pacing or I/R challenge. On the contrary, they exhibited a paradoxical enhancement in post-ischemic EP recovery compared to ND-fed controls. This improvement was associated with increased phosphorylation of canonical AMPK targets, including acetyl-CoA carboxylase (ACC) and raptor, consistent with the activation of a cardioprotective metabolic program. In sharp contrast, AMPK-deficient (AMPK-KD) hearts demonstrated heightened vulnerability to inducible ventricular tachycardia (VT), irrespective of diet. Conduction slowing emerged as an early EP abnormality in these hearts and served as the initial substrate (or ‘first hit’) that promoted their increased incidence of non-sustained VT. Notably, this conduction impairment arose in conjunction with an increase (rather than decrease) in Cx43 and Nav1.5 protein expression. Mechanistically, defective conduction in AMPK-KD hearts was linked to impaired autophagic degradation of intercalated disc proteins resulting from reduced phosphorylation of ULK1, a downstream effector of AMPK. Consequently, unphosphorylated Cx43 accumulated at the intercalated disc, likely replacing phosphorylated isoforms (p-Cx43). In addition, AMPK-KD hearts exhibited swollen, fragmented mitochondria and reduced levels of mitochondrial fusion proteins. Upon HFD challenge, this vulnerable mitochondrial substrate generated excessive reactive oxygen species (ROS) coinciding with accelerated repolarization. Together, impaired conduction and action potential shortening promoted VT sustenance in HFD-fed AMPK-deficient hearts.

**Conclusions:** Our findings identify AMPK as a key metabolic regulator that integrates redox balance, mitochondrial integrity, and protein homeostasis to preserve cardiac excitability during early nutrient overload. Loss of AMPK signaling, as occurs with aging and advanced metabolic disease, may therefore represent a pivotal mechanism linking HFD to increased SCD risk.

## INTRODUCTION

Obesity is a well-recognized and increasingly prevalent risk factor for cardiovascular disease and is strongly linked to a heightened risk of sudden cardiac death (SCD), primarily due to its association with ventricular arrhythmiasL(1, 2). In the United States alone, more than 100 million adults meet the clinical criteria for obesity, underscoring the urgent public health challenge posed by this conditionL(3). The cardiovascular burden of obesity extends beyond traditional risk factors, contributing significantly to the incidence of fatal arrhythmic events. A 2018 meta-analysis of 14 prospective cohort studies demonstrated that individuals with obesity have a two- to threefold higher risk of SCD compared to their normal-weight counterparts (4). This elevated risk has historically been attributed to the cumulative effects of chronic obesity on cardiac structure and function, including the development of coronary artery disease, myocardial hypertrophy, interstitial fibrosis, heart failure with preserved ejection fraction (HFpEF), and autonomic imbalanceL(1, 5). These pathophysiological changes promote a substrate that predisposes the heart to ventricular arrhythmias, often culminating in sudden cardiac arrest. However, emerging evidence suggests that arrhythmogenic remodeling may begin much earlier, during the initial phases of nutrient overload, well before overt obesity or structural cardiac changes become apparent. As such, there is growing interest in understanding how early metabolic disturbances triggered by high-fat diet (HFD) feeding contribute to electrical instability and arrhythmia vulnerability, independent of the long-term consequences of obesity *per se*.

Paradoxically, several clinical observations suggest that obesity may be associated with improved outcomes in certain cardiovascular contexts, a phenomenon referred to as the ‘obesity paradox’ (6, 7). While controversial, this concept raises the possibility that nutrient excess itself might engage adaptive mechanisms to confer protection (at least transiently) in a manner analogous to ischemic preconditioning.

AMP-activated protein kinase (AMPK) is a central regulator of energy homeostasis, mitochondrial integrity, and redox balance (8). In vitro studies have shown that fatty acids, such as palmitate, can stimulate AMPK through allosteric mechanisms (9). Furthermore, pharmacological activation of AMPK *in vivo* is known to preserve myocardial viability, limit infarct size and improve mitochondrial function following ischemia-reperfusion (I/R) injury (10, 11). While the role of AMPK in mediating protective signaling in response to energetic and metabolic stress programs is well established, its function in regulating cardiac electrophysiological (EP) responses under these conditions remains completely unknown. We hypothesized that early nutrient overload intrinsically activates AMPK as an adaptive mechanism to protect against arrhythmogenesis, and that loss of AMPK signaling predisposes the heart to adverse EP remodeling, particularly under HFD conditions.

We demonstrate that WT hearts exposed to short-term HFD exhibit paradoxical protection against arrhythmias, characterized by enhanced post-ischemic EP recovery and increased phosphorylation of canonical AMPK targets, consistent with activation of a cardioprotective metabolic program. In contrast, AMPK-KD hearts display baseline conduction slowing, driven by impaired ULK1-mediated autophagy. This defect results in the accumulation of dephosphorylated, inactive Cx43 at the intercalated disc, disrupting gap junction coupling. This conduction abnormality constitutes the first electrophysiological “hit” and is sufficient to increase the incidence of non-sustained VT. Concomitantly, AMPK-KD hearts exhibit mitochondrial ultrastructural abnormalities, reflecting a second layer of vulnerability. Upon HFD challenge, this compromised mitochondrial network becomes a source of excessive ROS which precipitate sustained VT. Together, these findings support a two-hit model of arrhythmogenesis in AMPK-deficient hearts and establish AMPK as a key regulator of cardiac electrical stability during early metabolic stress.

## RESULTS

In this study, we sought to determine how early nutrient overload, specifically from dietary fat, impacts EP remodeling prior to the development of overt obesity, independent of chronic obesity-related confounders. To do this, we employed a model of short-term HFD-induced metabolic stress. Male and female C57BL/6N wild-type (WT) mice were fed either a normal diet (ND) or a 45% HFD for 8 weeks, starting at 4 weeks of age (Figure 1A). By the end of the feeding period, HFD-fed mice exhibited only modest weight gain (∼5 g) compared to ND controls, accompanied by a proportional increase in heart weight. As a result, the heart weight-to-body weight ratio remained unchanged (p = 0.8) (Figure 1B, C). Plasma glucose and triglyceride levels were also unaltered (Figure 1D), and echocardiographic analysis revealed no differences in systolic or diastolic function between groups (Figure 1E). These data confirm that short-term HFD feeding does not induce the overt metabolic or hemodynamic abnormalities typically associated with advanced obesity.

**Figure 1:**
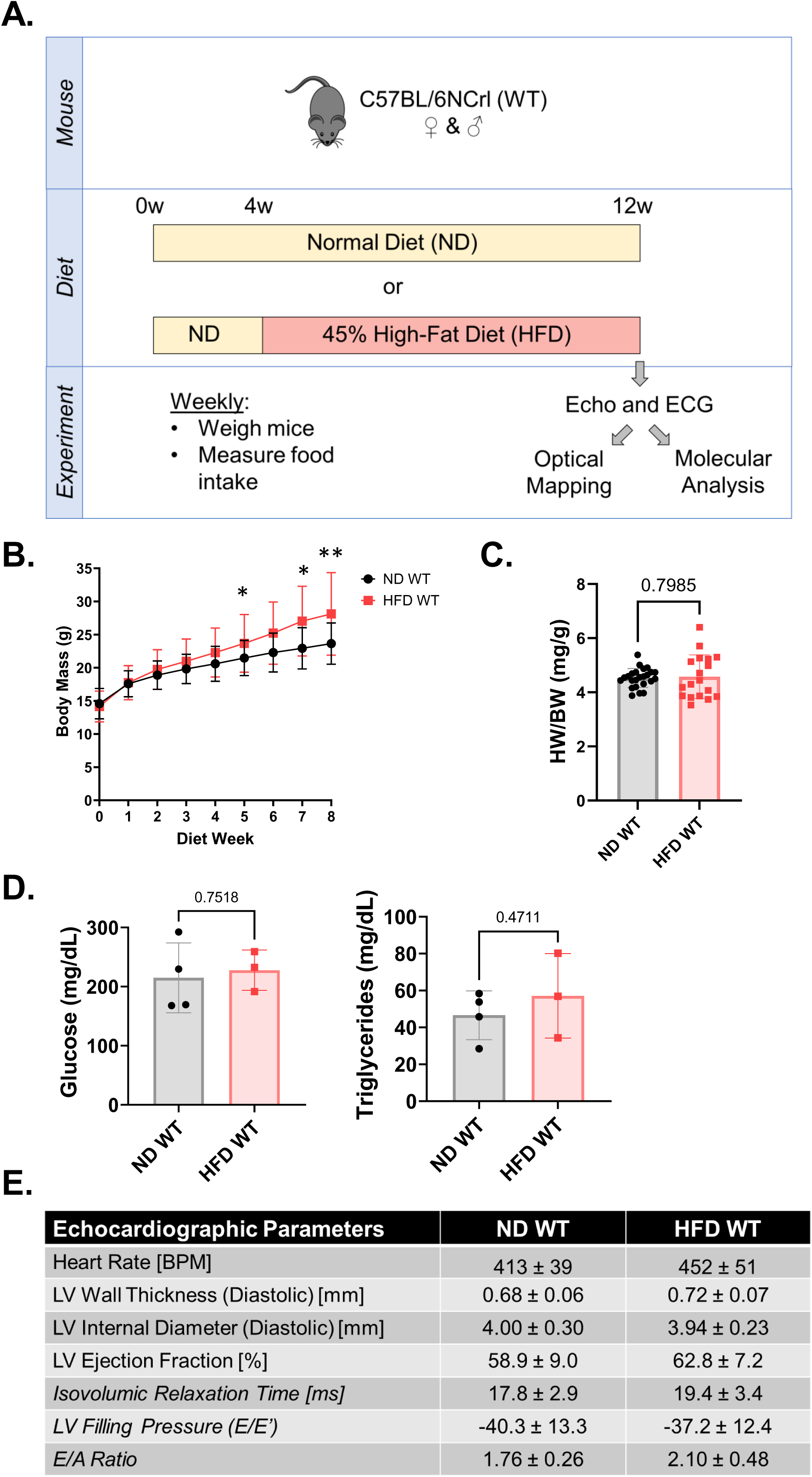
Establishing a model of high fat diet induced metabolic stress. **A.** Experimental model and timeline. Four-week-old mice of either sex are placed on normal or high fat (45%) diets for 8 weeks. At the terminal (12 weeks of age), mice undergo echocardiography and surface ECG recordings and are sacrificed for detailed electrophysiological studies using high-resolution optical action potential mapping or molecular and histological analyses. **B.** Average body mass during the 8-week regimen of normal (ND; n=17) or high fat (HFD; n=31) diet. **C.** Average heart weight normalized to total body weight (HW/BW) in ND vs HFD groups. **D.** Plasma levels of glucose (left) and triglycerides (right) after 8 weeks of ND and HFD (n=3 per group). **E.** Echocardiographic parameters after 8 weeks of ND (n=11) or HFD (n=10). No significant differences were found between the two groups. * indicates p <0.05, ** indicates p< 0.01

A central goal of this study was to assess whether early nutrient stress of high fat, prior to significant weight gain or left ventricular (LV) dysfunction modulates EP function and arrhythmia susceptibility. To test this, we subjected isolated hearts from ND- and HFD-fed mice to two stress protocols. First, rapid pacing up to 25 Hz, a stress protocol that is known to induce calcium overload related electrical instability, failed to provoke arrhythmias in either group (Figure 2A, B). Next, hearts underwent 12 minutes of global no-flow ischemia followed by reperfusion. Strikingly, HFD-fed WT hearts not only lacked reperfusion-induced arrhythmias but demonstrated significantly improved post-ischemic EP recovery. Shown in Figure 2C are representative depolarization isochrones from ND- and HFD-fed hearts at baseline (pre-ischemia) and following 12 minutes of reperfusion, illustrating full restoration of conduction to pre-ischemic levels in HFD hearts but not in ND controls. Notably, 100% of HFD hearts exhibited near-complete recovery of pre-ischemic conduction, defined as normalized conduction velocity (CV) exceeding 90% of baseline, whereas only 1 of 6 ND hearts met this threshold (p = 0.0152, Figure 2C, right). Action potential (AP) amplitude also recovered more robustly in HFD-fed hearts (Figure 2D). These findings suggest that short-term HFD exposure activates a cardioprotective program that enhances recovery from ischemic stress, even in the absence of obesity or LV dysfunction. Based on previous reports that exposure to fatty acids at least *in vitro* can allosterically activate AMPK via the conserved β-subunit ATM site, we hypothesized that *in vivo* AMPK activation by HFD underlies this effect (9). Supporting this, phosphorylation levels of the canonical AMPK targets acetyl-CoA carboxylase (ACC) and raptor were significantly increased in HFD-fed WT hearts, despite no change in total AMPK levels (Figure 3A, B). Phosphorylated-to-total protein ratios increased 2- to 3-fold, indicative of enhanced AMPK activity.

**Figure 2:**
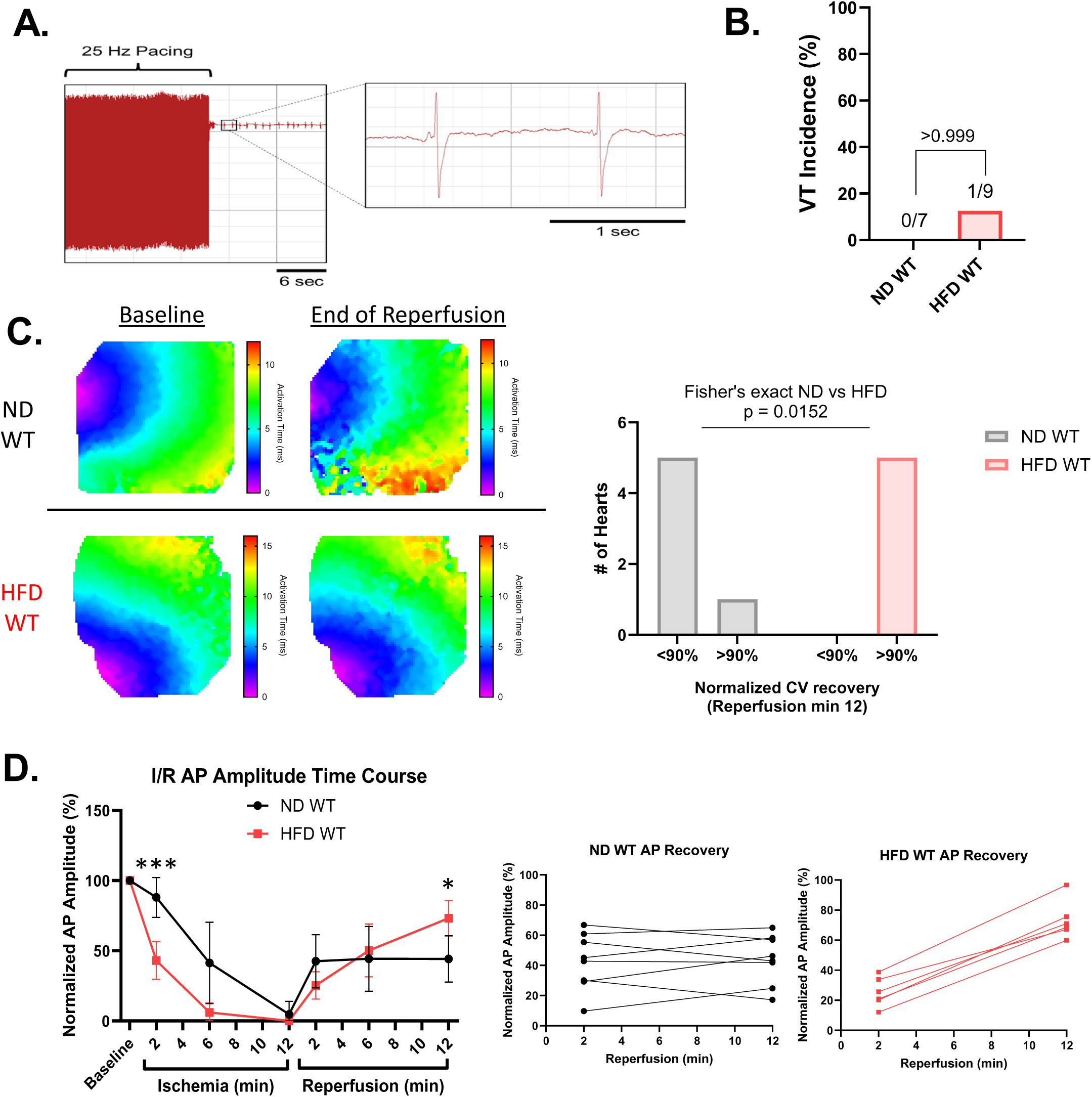
Response to arrhythmia provocation protocols of burst pacing and ischemia-reperfusion suggest activation of cardioprotective signaling in HFD hearts. **A.** Representative ex vivo volume conducted ECG trace from VT negative ND heart showing typical response to burst stimulation with rapid return to normal rhythm. **B.** Rapid pacing VT incidence rates in ND (0%) and HFD (11.1%) groups (Fisher’s exact p>0.999). VT incidence was based on episodes lasting for at least 2 minutes. **C.** Quantification of post-ischemic recovery of CV at minute 12 of reperfusion. CV is normalized to pre-ischemic CV for each heart and recovery is separated into <90% (residual damage) and >90% (full recovery). ND WT (n=6) and HFD WT (n=5) are compared with a Fisher’s exact test (p =.0152) Representative maps for ND WT and HFD WT at baseline (pre-ischemia) and reperfusion minute 12 are shown. **D.** Average AP amplitude during the course of I/R normalized to basal pre-ischemic levels in WT (black) and HFD (red) hearts. Slope of recovery during reperfusion showing increasing values in HFD but not ND hearts. * indicates p <0.05, ** indicates p< 0.01, *** indicates p< 0.001

**Figure 3:**
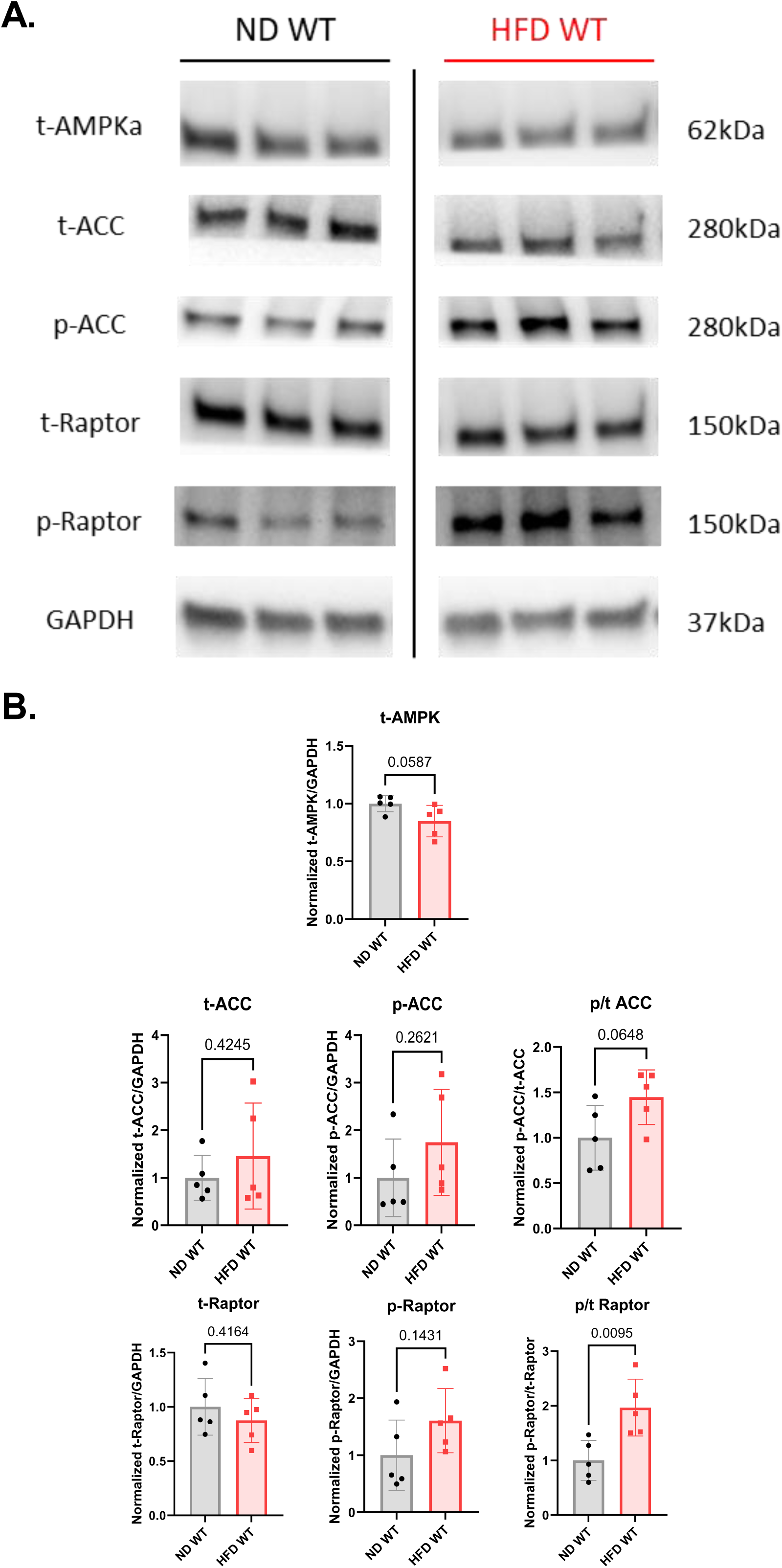
HFD enhances AMPK activity in WT hearts, supporting its involvement in cardioprotective mechanisms. **A.** Representative western blots of total AMPK, total ACC, phospho-ACC, total Raptor, phospho-raptor and GAPDH in ND and HFD WT hearts. **B.** Quantifications of total AMPK, total ACC, phospho-ACC, total Raptor, phospho-raptor relative to GAPDH, and phosphorylation to total ratios of ACC and Raptor.

To determine whether this putative protective response persists with prolonged metabolic stress, we repeated the I/R challenge in hearts from mice fed HFD for 16 weeks. As expected, these mice exhibited substantial weight gain. Unlike the 8-week HFD group, 16-week HFD hearts displayed marked reductions in AMPK signaling, including diminished phosphorylation of AMPK and its downstream targets (Supplemental Figure 1). Functionally, these hearts failed to restore conduction following reperfusion (Supplemental Figure 1) and were partially susceptible to pacing-induced VT (40% induction rate).

To directly assess whether AMPK is required for EP protection against early metabolic stress in 8-week HFD fed mice, we used AMPK kinase-dead (AMPK-KD) mice, which express a catalytically inactive AMPKα2 isoform in the heart and skeletal muscle (Figure 4A) (12). In stark contrast to WT mice, AMPK-KD hearts failed to restore conduction following 8 weeks of HFD feeding and exhibited worsened post-ischemic conduction recovery (Figure 4B, C). Even under ND conditions, AMPK-KD hearts showed increased VT susceptibility, although the episodes were much shorter in duration than those seen with HFD (Figure 4D, E). These findings support a model in which AMPK deficiency irrespective of diet establishes a vulnerable substrate that promotes the onset of VT.

**Figure 4:**
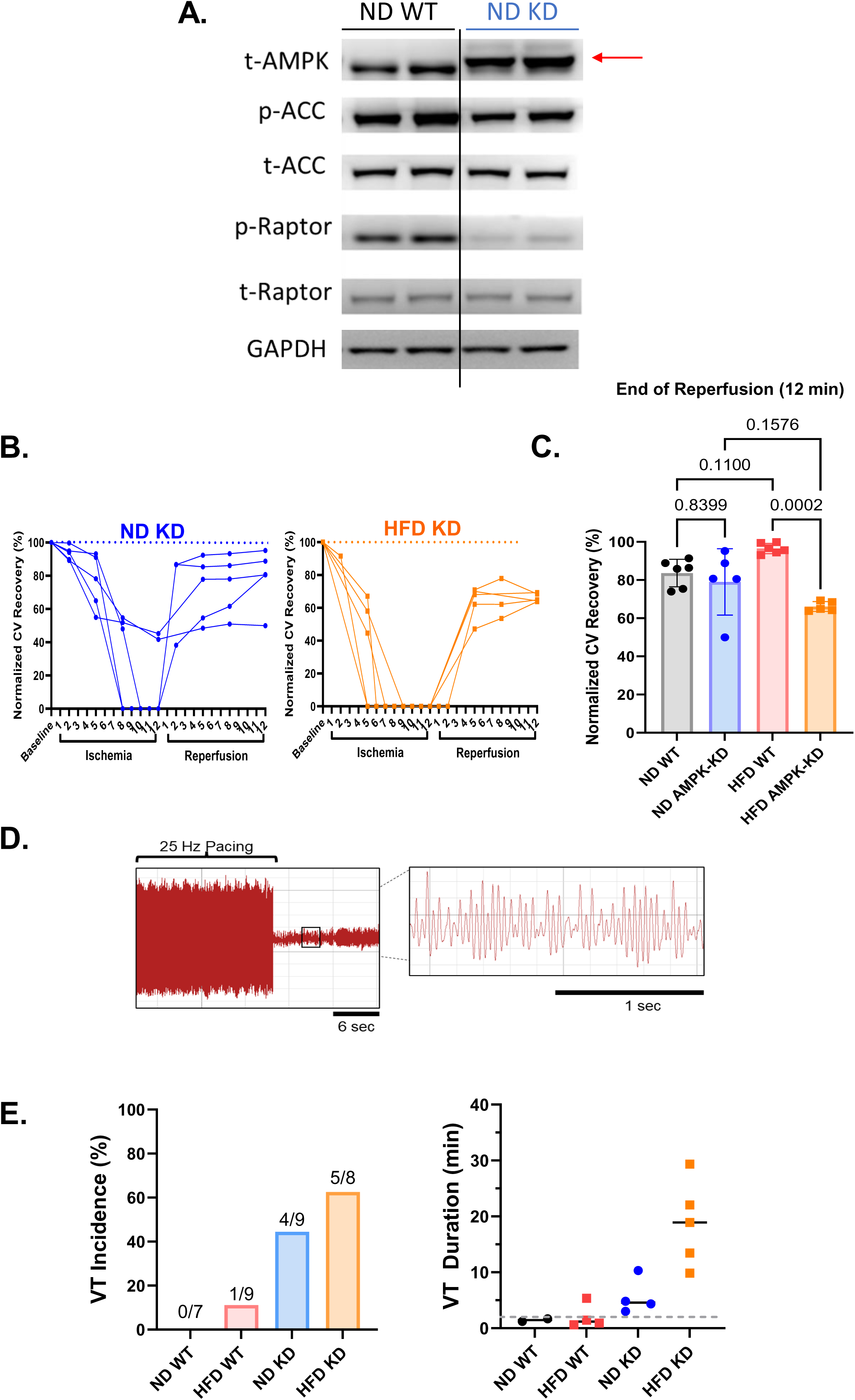
Genetic inactivation of cardiac AMPK leaves HFD challenged hearts susceptible to marked worse I/R recovery and severe arrhythmias. **A.** Western blots of AMPK and its downstream signaling targets in hearts from WT and AMPK kinase dead (KD) mice expressing a c-myc-tagged K45R alpha isoform of AMPK with defective downstream signaling. The c-myc-tagged K45R mutant isoform migrates to a higher band compared to the native AMPK isoform in WT hearts (red arrow). **B.** Changes in myocardial conduction velocity (CV) during the course of global 12-minute no-flow ischemia followed by reperfusion, normalized in each heart to basal pre-ischemic CV in multiple ND (left) and HFD (right) KD hearts. **C.** Quantification of the extent of post-ischemic recovery measured at the end of the 12-min reperfusion phase normalized relative to baseline pre-ischemic levels in each heart as its own control. **D.** Representative ECG of a sustained VT episode following rapid pacing challenge in ND KD heart. **E.** Rates of sustained VT incidence defined as episodes lasting for at least 2 min (left) and extent of arrhythmia severity indexed by VT episode duration (right) in each group. Dashed line indicating 2 min cutoff for sustained VT.

To investigate the basis of this vulnerability, we first examined the basal EP properties of AMPK-KD hearts under ND conditions. While AP duration was similar to WT (Figure 5A), AMPK-KD hearts exhibited an average 30% reduction in CV (Figure 5B) and increased incidence of anomalous conduction patterns with areas of conduction block (Figure 5D, E) despite the absence of fibrosis or structural remodeling (Figure 5C). Western blotting revealed paradoxical upregulation, not downregulation, of Nav1.5 and connexin-43 (Cx43), suggesting that altered overall expression was not responsible for conduction slowing (Figure 6A, B).

**Figure 5:**
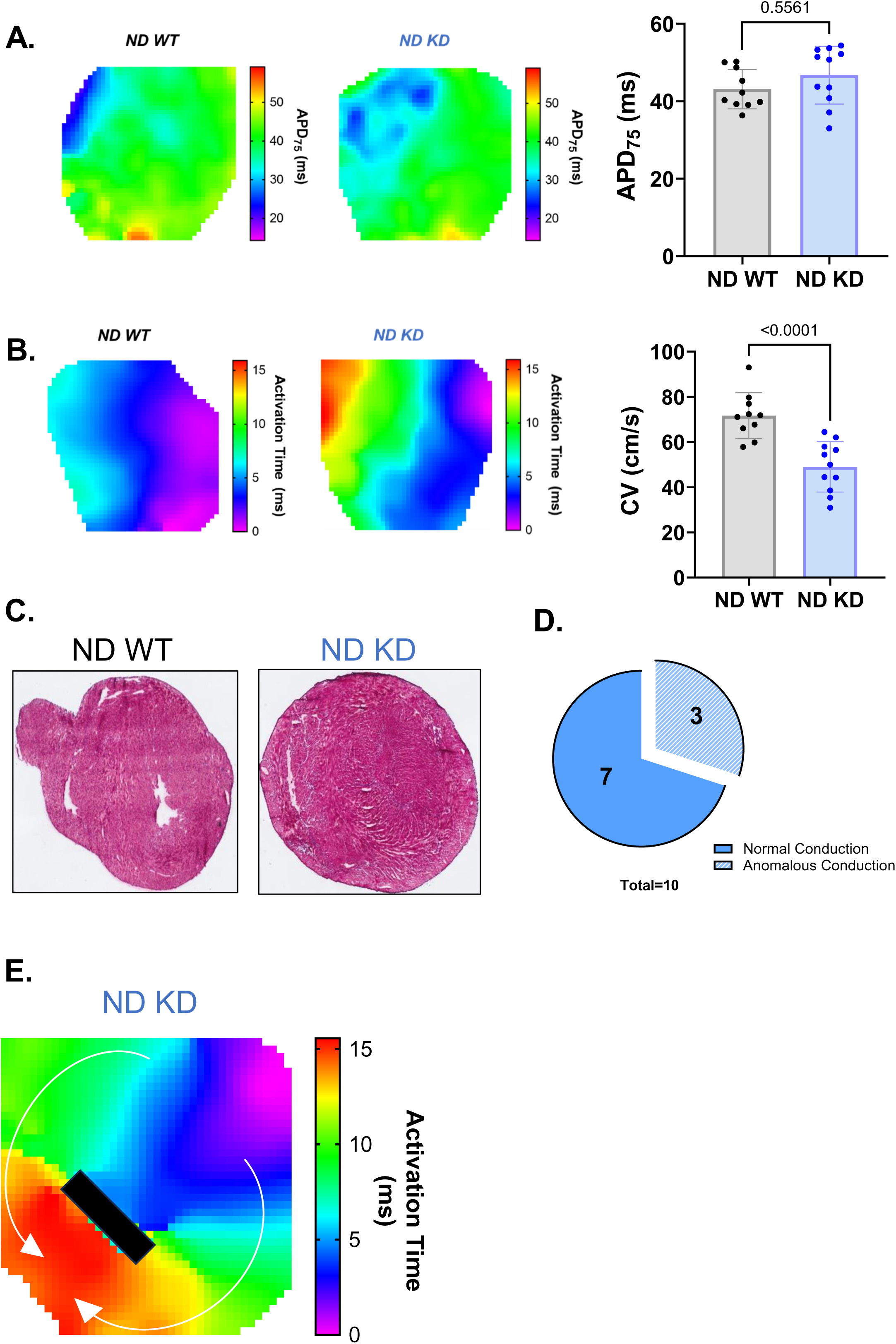
Inactivation of AMPK leads to slowed CV and aberrant conduction in AMPK-kinase dead mice. **A.** Representative contour maps recorded at PCL 100ms in WT (left) and KD (right) hearts showing action potential duration. Quantification of average APD75 at baseline for ND WT and KD groups shown beside. **B.** Representative depolarization isochrone maps recorded at PCL 100ms in WT (left) and KD (right) hearts. Quantification of average CV at baseline for ND WT and KD groups shown beside. **C.** Representative Masson’s trichrome stains in ND WT and ND KD groups revealing normal structure with no evidence of fibrosis. **D.** Chart indicating the percentage of KD hearts which exhibit an anomalous pattern of epicardial activation at PCL 100ms (n=10). **E.** Activation activation map showing anomalous conduction pattern with functional line of block (black rectangle, arrows indicate conduction route).

**Figure 6:**
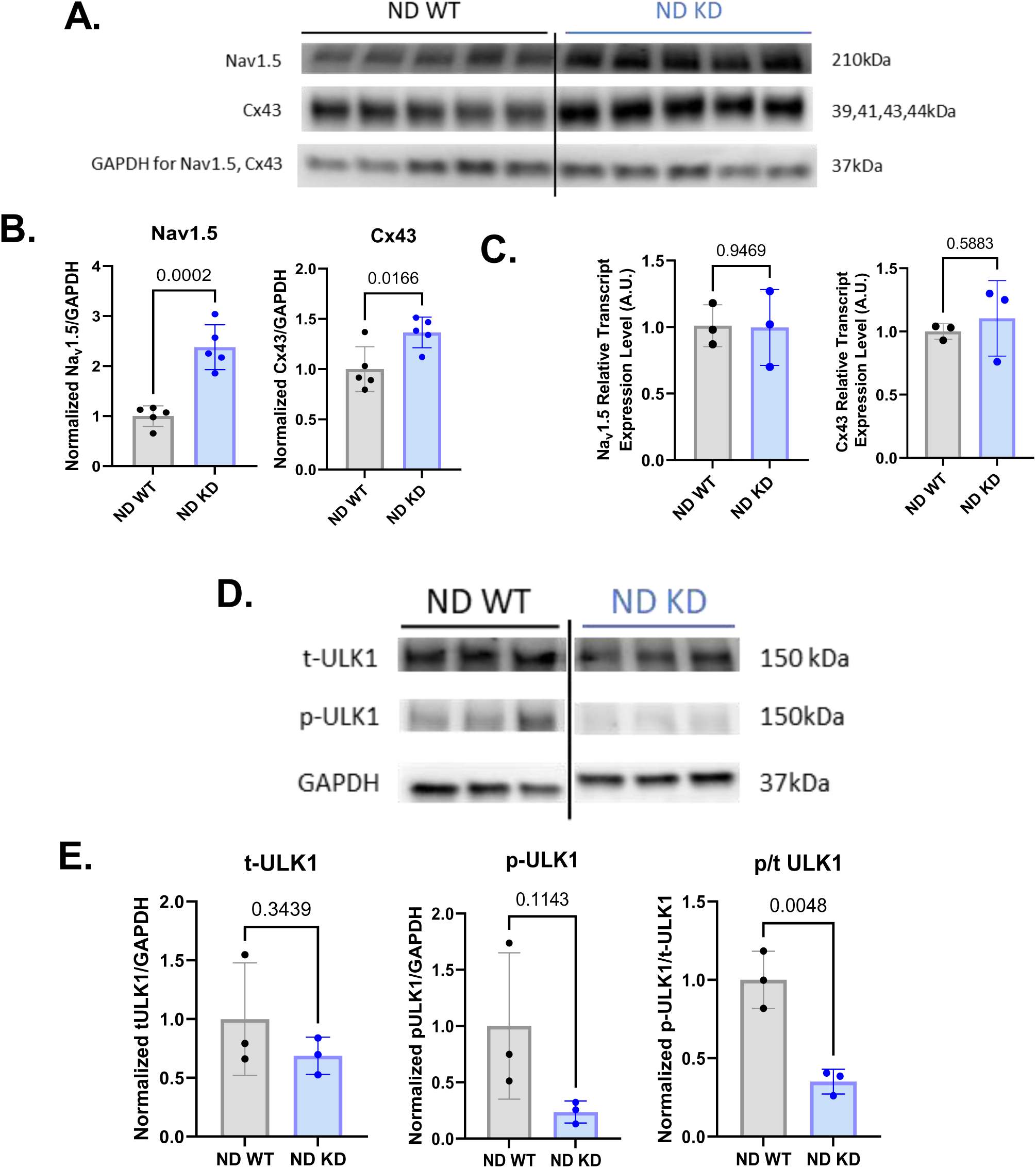
Molecular mechanism of conduction defects caused by AMPK pathway inactivation. **A.** Western blot showing conduction related proteins Nav1.5, Cx43, and GAPDH in ND WT and ND KD groups (n=5 per group). **B.** Quantification of A. **C.** qPCR results of Nav1.5 and Cx43 show similar transcript levels between groups (n=3 per group). **D.** Western blot shows decreased phosphorylation of ULK1 (n=3 per group). **E.** Quantification of D showing marked reduction in a key AMPK regulated autophagy pathway (p-ULK1/tULK1).

Given the established role of AMPK in regulating autophagy and its enriched localization at intercalated discs in the phosphorylated state, a pattern that we confirmed, we hypothesized that AMPK plays a key role in the homeostatic turnover of intercalated disc proteins that are critical for cardiac conduction namely Cx43, which has a short half-life (∼90 minutes) (14). In AMPK-KD hearts, normal transcript levels (Figure 6C) combined with impaired autophagic degradation may allow accumulation of dysfunctional Cx43 isoforms at mechanically stressed junctions. Supporting this, phosphorylation of ULK1, a key AMPK target involved in autophagy initiation, was significantly reduced in AMPK-KD hearts (Figure 6D, E). Confocal imaging showed reduced colocalization of phosphorylated Cx43 (p-Cx43) at Ser-368, but not total Cx43, with the intercalated disc marker N-cadherin (Figure 7) consistent with accumulation of the dephosphorylated isoform at gap junction loci.

**Figure 7:**
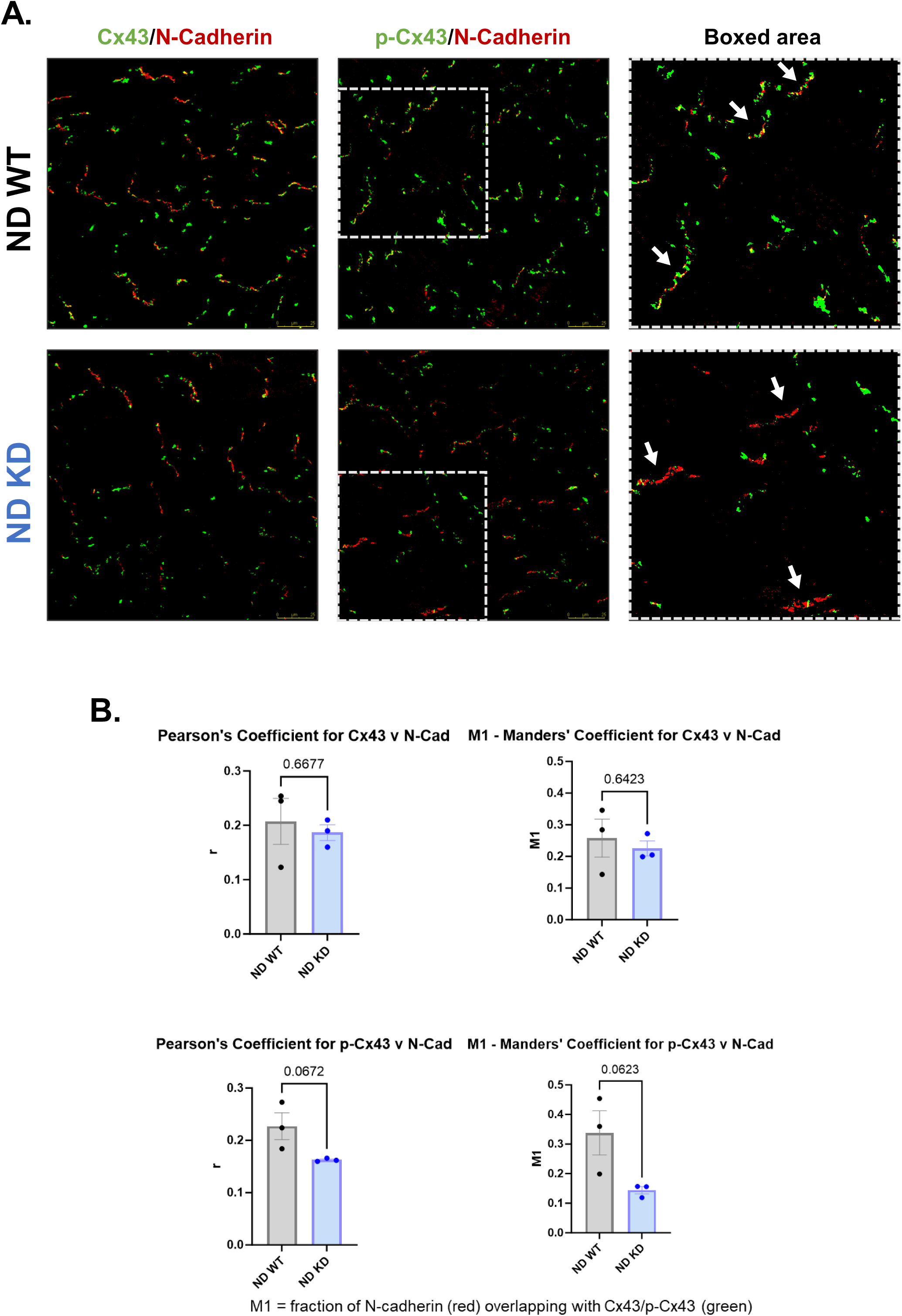
Phosphorylated Cx43 downregulated at intercalated disk when AMPK is inactivated. **A.** Representative confocal images showing the localization of the intercalated disc protein, N-Cadherin (red), and Cx43 (green, left) or p-Cx43 (green, middle) at 40x. Boxed areas of p-Cx43 images are shown on the right for visual clarity. White arrows indicate intercalated disks. **B.** Quantification of the co-localization of N-Cadherin/Cx43 (top) and N-Cadherin/p-Cx43 (bottom) by Pearson’s coefficient and Mander’s M1 coefficient. Data reveal major reduction in colocalization between p-Cx43 and N-Cadherin in the ND KD relative to ND WT group.

Given the contrasting EP responses to ischemia between HFD-fed WT and AMPK-KD hearts, and AMPK’s established role in regulating mitochondrial fission and fusion events, we next examined mitochondrial structure, dynamics and related protein expression. Transmission electron microscopy revealed enlarged, swollen mitochondria in AMPK-KD hearts, despite preserved mitochondrial density and aspect ratio (Figure 8A). Western blotting showed increased DRP1 expression, decreased phosphorylation at the AMPK-regulated inhibitory Ser-637 site, and reduced levels of fusion proteins MFN1 and OPA1 (Figure 8B, C). These findings are consistent with a shift toward mitochondrial fission and impaired mitochondrial homeostasis.

**Figure 8:**
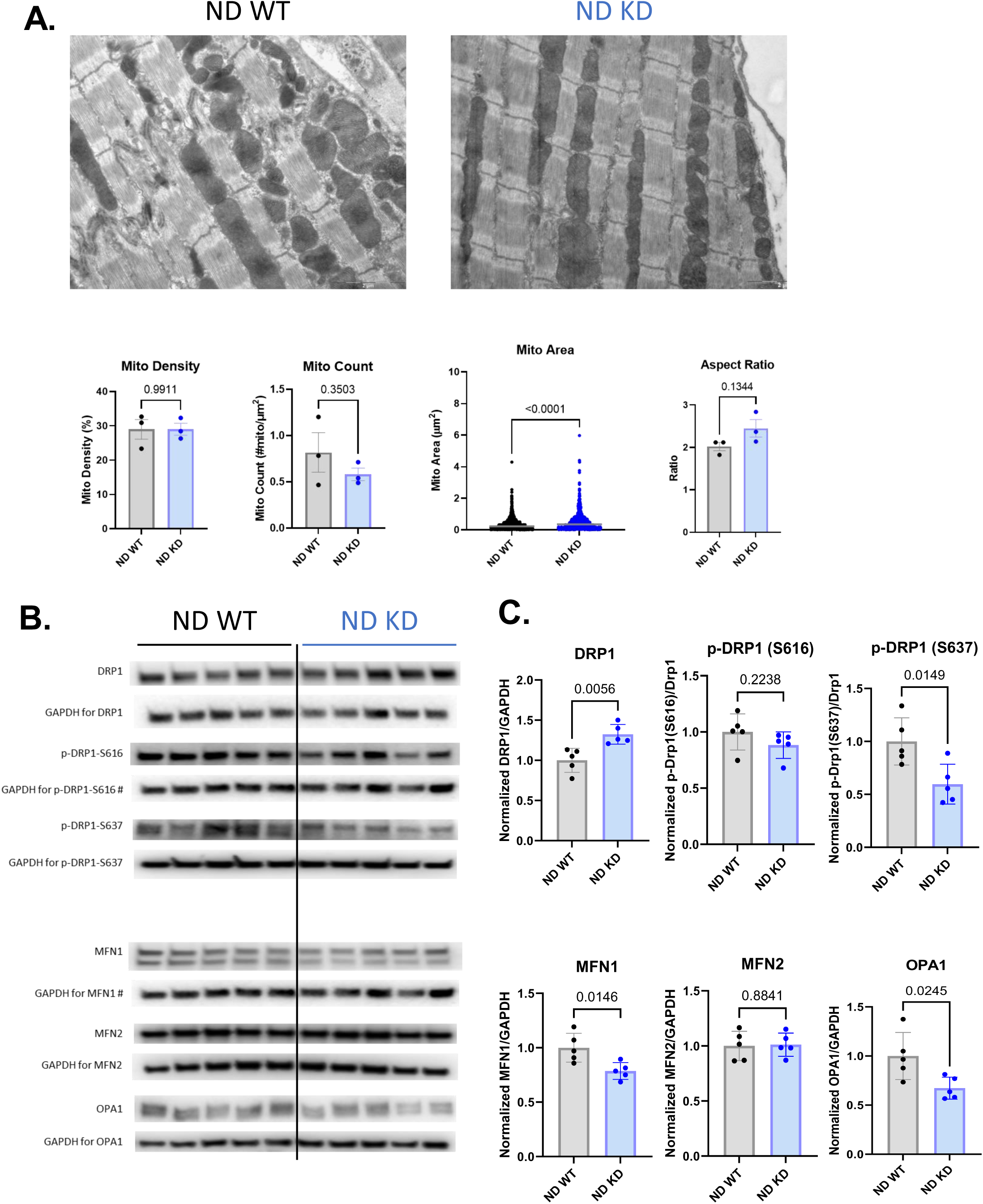
Loss of AMPK activity disrupts the AMPK-DRP1 axis resulting in abnormal mitochondrial ultrastructure. **A.** Electron microscopy images of ventricular tissue from ND WT and ND KD mice. Quantification of mitochondrial density, count, area, and aspect ratio is also shown under the representative EM images for each group. **B.** Western blots of key mitochondrial dynamics proteins namely DRP1, phosphorylated (p) DRP1 at S616, p-DRP1 at S637, MFN1, MFN2, and OPA1 (n=5 per group). **C.** Quantification of mitochondrial dynamics protein expression.

Lastly, we evaluated whether mitochondrial defects in AMPK-deficient hearts promote oxidative stress and contribute to electrical instability. Dihydroethidium (DHE) staining revealed significantly elevated reactive oxygen species (ROS) levels in HFD-fed AMPK-KD hearts compared to all other groups (Figure 9A). This pro-oxidant state was associated with accelerated repolarization, evidenced by significant shortening of the AP duration, seen exclusively in HFD-fed AMPK-KD hearts (Figure 9C, D). This arrhythmogenic EP profile consisting of impaired conduction combined with accelerated repolarization, likely underlies the heightened susceptibility to sustained VT in this group (Figure 9E), implicating ROS as a key mechanism by which AMPK deficiency underlies the sustenance of malignant VT during early nutrient overload.

**Figure 9:**
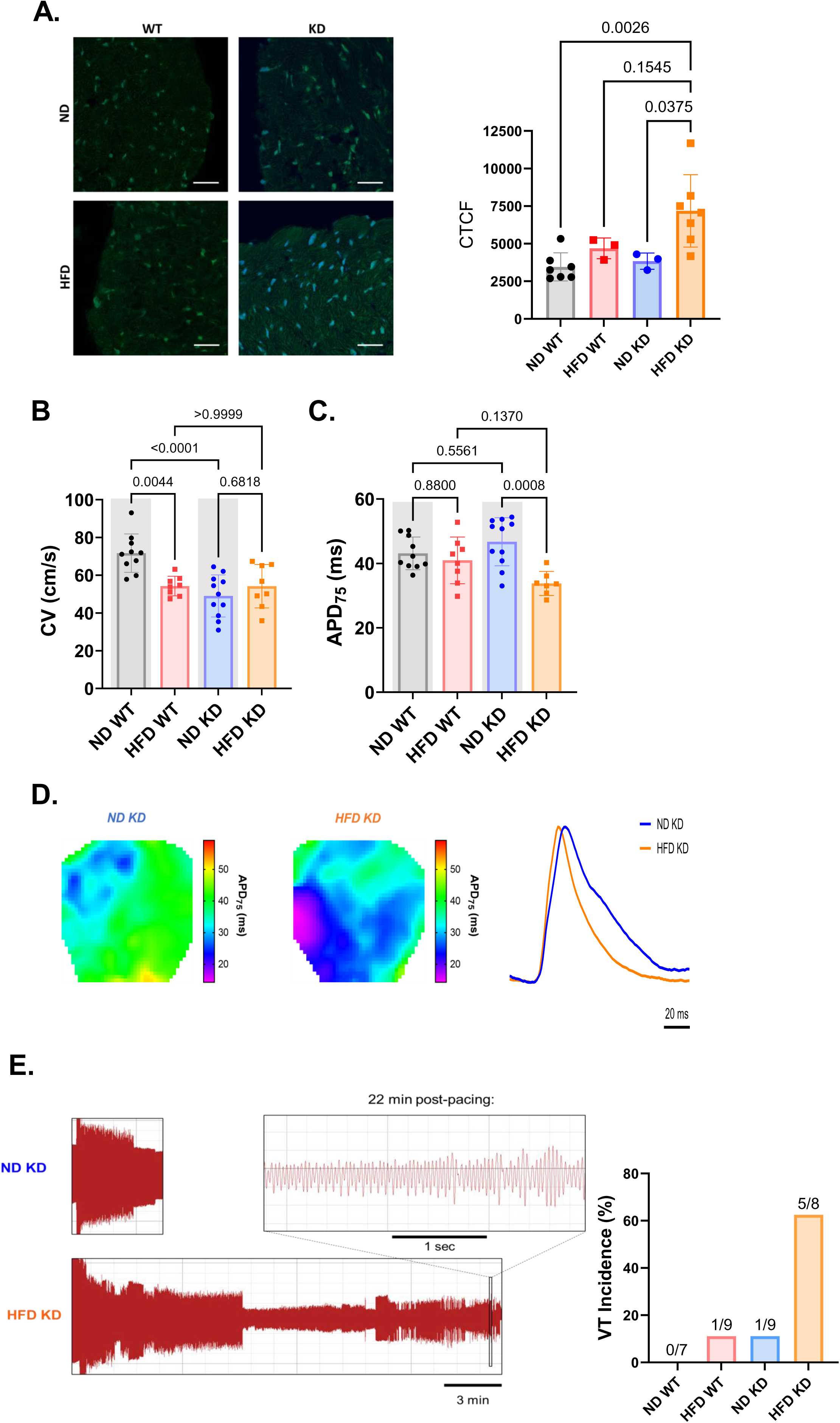
Excessive ROS generation from HFD in KD background drives APD shortening as a second hit increasing susceptibility to severe VT episodes. **A.** Average basal ROS levels in each group quantified from DHE staining. Representative images of each group shown on the left. **B.** Average CV and **C.** average APD75 at PCL100. Gray areas indicate the data were previously shown in Figure 5 and are reshown here for visual clarity. **D.** Representative APD contour maps from ND KD and HFD KD groups showing shortening of the APD (darker colors) and representative action potential traces showing abrupt acceleration of repolarization in the HFD KD group (orange trace) relative to ND KD (blue). **E.** Representative ECGs from ND KD (top) and HFD KD (bottom) VT episodes showing increased severity in the HFD group. When severe VT is defined as sustained past 5 minutes, HFD KD has the highest incidence of over 60%.

## DISCUSSION

Our study identifies AMPK as a critical regulator of cardiac electrophysiological stability during early HFD-induced metabolic stress. We demonstrate that short-term HFD exposure, prior to the onset of obesity, hyperglycemia, or structural remodeling, paradoxically enhances post-ischemic EP recovery in WT hearts. This protection is associated with increased phosphorylation of canonical AMPK targets, consistent with the activation of a cardioprotective metabolic program.

In contrast, AMPK-deficient (AMPK KD) hearts fail to engage this adaptive response and exhibit a baseline susceptibility to inducible arrhythmias, independent of diet. This vulnerability stems from an intrinsic EP defect characterized by slowed, and in some cases aberrant, conduction. This conduction abnormality arises from impaired ULK1-dependent autophagy, which specifically affects intercalated disc proteins, a region where AMPK is normally enriched. Loss of AMPK at this structurally and functionally critical site disrupts proteostatic control, leading to the accumulation of hypophosphorylated Cx43 at the intercalated disc.

Alongside these electrical abnormalities, AMPK KD hearts exhibit mitochondrial ultrastructural defects, including swelling, fragmentation, and downregulation of the fusion proteins MFN1 and OPA1. When subjected to the same HFD regimen that elicits protection in WT hearts, these structurally compromised mitochondria in AMPK KD hearts generate excessive reactive ROS, potentially introducing a secondary electrophysiological insult in the form of APD shortening. The convergence of conduction slowing and accelerated repolarization culminates in sustained ventricular arrhythmias, which are markedly prolonged in duration in AMPK KD hearts. These findings support a two-hit model of arrhythmogenesis under impaired AMPK signaling. A primary defect in conduction predisposes the myocardium to arrhythmia initiation, while a secondary insult from mitochondrial ROS further destabilizes repolarization to sustain VT.

Notably, conduction remodeling in AMPK deficient hearts does not align with classic histopathologic substrates such as fibrosis or downregulation of major conduction-related proteins. On the contrary, alpha subunits of Nav1.5 and Cx43 are paradoxically upregulated. Rather, the phenotype is defined by reduced ULK1 phosphorylation, reinforcing AMPK’s role in initiating autophagy at the intercalated disc and maintaining proteostasis of key conduction proteins that are expressed at this cellular locus. Previous studies have shown that Cx43 phosphorylation is required for its incorporation into autophagosomes and subsequent degradation (15). Our earlier work in a canine model of pacing-induced heart failure demonstrated that Cx43 dephosphorylation at the intercalated disc correlates with conduction slowing, independent of total expression levels (16). Building on these findings, the current study identifies AMPK as an upstream regulator of Cx43 phosphorylation, turnover, and localization, linking for the first time metabolic stress signaling to functional gap junction integrity and remodeling.

Finally, our earlier work showed that administration of the fatty acid palmitate preserves mitochondrial energetic and redox balance in relatively young (2-3 month) db/db mice, protecting against mechanical dysfunction despite underlying diabetic stress (17). We subsequently found that db/db mouse hearts at this stage maintained normal mitochondrial network ultrastructure (18). These findings support the notion that dietary fat can engage cardioprotective mechanisms when mitochondrial architecture remains intact, consistent with our present results in wild-type hearts exposed to short-term HFD. Together, they reinforce the concept that mitochondrial substrate quality and AMPK signaling status dictate whether nutrient excess triggers cardioprotective adaptation or maladaptive remodeling. Importantly, mitochondrial disruption in AMPK KD hearts further amplifies electrical instability to result in excessively prolonged VT episodes. These hearts exhibit increased DRP1-mediated fission and reduced mitochondrial fusion, promoting ROS overproduction during metabolic stress. This oxidative stress likely accelerates repolarization and acts synergistically with baseline conduction defects to create a highly arrhythmogenic substrate that is suitable not only for VT initiation but also maintenance of reentrant activation.

Collectively, these data offer a mechanistic framework to reinterpret the so-called obesity paradox. While obesity is broadly recognized as a risk factor for SCD, several studies have paradoxically associated obesity with improved outcomes in certain cardiovascular settings. Our findings suggest that this paradox may reflect not obesity per se, but rather the heart’s capacity to activate protective metabolic stress responses at least transiently. In WT hearts, preserved AMPK signaling facilitates adaptive remodeling in response to early nutrient excess, maintaining electrical stability. In contrast, AMPK deficiency, whether due to genetic depletion, aging, or advanced metabolic disease, permits the same stressor to unmask a latent arrhythmogenic phenotype.

This study expands AMPK’s role beyond energy homeostasis and infarct limitation to include dynamic regulation of cardiac conduction through mitochondrial quality control and gap junction proteostasis. Given the rapid turnover of Cx43 and its reliance on post-translational regulation, it serves as a sensitive target through which AMPK modulates cell-to-cell electrical coupling. Mislocalization of phosphorylated Cx43 in AMPK KD hearts exemplifies how impaired stress signaling leads to impaired conduction.

Finally, systemic conditions known to impair AMPK activity, such as aging, insulin resistance, and type 2 diabetes, are also strongly associated with increased arrhythmic risk (Coughlan, 2013; Reznick, 2007). Our findings suggest that therapeutic strategies aimed at restoring or enhancing AMPK activity may mitigate arrhythmic vulnerability by preserving mitochondrial and intercalated disc homeostasis under metabolic stress.

### Limitations

Several limitations warrant acknowledgment. First, all experiments were conducted in C57BL/6N mice, which may limit generalizability due to potential strain-specific differences in metabolic adaptation and arrhythmia susceptibility. Second, although we extensively characterized mitochondrial morphology and redox status in AMPK KD hearts, we did not assess whether HFD alters mitochondrial structure or function in WT hearts. Future studies are needed to determine whether mitochondrial protection in WT hearts is transient and reversible. Third, optical mapping was performed under electromechanical uncoupling with blebbistatin, which may affect calcium dynamics and restitution properties. Fourth, although we implicate impaired ULK1 signaling in defective autophagy and gap junction remodeling, we did not directly measure autophagic flux or utilize ULK1-specific knockout models. The precise downstream mediators linking AMPK to Cx43 turnover remain to be elucidated. Lastly, while our results support a two-hit mechanism involving conduction slowing and ROS-driven repolarization defects, this model would benefit from targeted rescue experiments. These may include pharmacologic activation of autophagy or mitochondrial antioxidants to dissect the relative contributions of each insult to the observed arrhythmic phenotype.

### Conclusions

Our findings position AMPK as a central metabolic safeguard that integrates mitochondrial quality control, redox balance, and protein turnover at the intercalated disc to preserve cardiac excitability during early nutrient stress. In wild-type hearts, short-term HFD exposure activates an AMPK-dependent protective program that enhances recovery from ischemic injury. In contrast, AMPK deficiency unmasks a latent arrhythmogenic phenotype via a two-hit mechanism: conduction slowing from impaired Cx43 turnover and mislocalization, followed by HFD-induced ROS generation and repolarization acceleration.

These insights provide a mechanistic explanation for how early nutrient excess produces divergent electrophysiological outcomes depending on AMPK status. They also offer a reinterpretation of the obesity paradox as an HFD paradox, wherein the capacity for adaptive AMPK signaling dictates vulnerability to sudden cardiac death. Our findings highlight AMPK as a promising therapeutic target for arrhythmia prevention in aging, insulin resistance, and cardiometabolic disease.

## METHODS

### Animal Procedures

All animal procedures were approved by the Yale Institutional Animal Care and Use Committee (IACUC) and carried out by trained personnel. AMPK kinase-dead (KD) mice, carrying a dominant-negative K45R mutation in cardiac and skeletal muscle, along with their wild-type (WT) littermates, were used for all experiments (12). At 4 weeks of age, mice were randomly assigned to either a normal diet (ND; 18 kcal% fat, Inotiv, #2018S) or a high-fat diet (HFD; 45 kcal% fat, Research Diets, #D12451) for 8 weeks. Mice were fed ad libitum and maintained on a 12-hour light/dark cycle. Weight and food intake were monitored weekly. Both male and female mice were included and analyzed together. To confirm the absence of overt obesity, plasma glucose and triglyceride levels were measured.

At 12 weeks of age, mice underwent echocardiography, optical mapping, or tissue collection. Mice were injected intraperitoneally with 0.3 mL of a 1:1 heparin:sodium chloride solution and euthanized 20 minutes later by cervical dislocation. Hearts were rapidly excised, and plasma was collected from the chest cavity for analysis.

### Echocardiography

Mice were anesthetized with 1–2% isoflurane and transthoracic echocardiography was performed using a Vevo 2100 high-resolution ultrasound system, similar to previously described.(19) Body temperature was maintained throughout.

### Optical Mapping

Optical mapping was performed as previously described (20). Briefly, 12-week-old mice were injected with 0.3 mL of a 1:1 heparin:saline solution, euthanized, and hearts were excised and cannulated on a Langendorff apparatus (Fisnar, #8001288). Hearts were perfused with Tyrode’s solution (129 mmol/L NaCl, 24 mmol/L NaHCO3, 4 mmol/L KCl, 1 mmol/L MgCl2, 11.2 mmol/L glucose, 1.8 mmol/L CaCl2, 1.2 mmol/L KH2PO4) and submerged in a bath of oxygenated Tyrode’s. Blebbistatin (10 µM, Sigma-Aldrich, #B0560) was added to suppress motion artifacts. Perfusion pressure was maintained at ∼70 ± 15 mmHg by adjusting flow (2.0 ± 1 mL/min) and temperature was held at 37 ± 1°C.

Hearts were loaded with 30 µL of 2.5 mM Di-4-ANNEPS (ThermoFisher #D1199) via inline injection. A pacing electrode was placed in the left ventricular apex. ECG and high-resolution optical signals were acquired as previously described with a 6×6 mm² field of view (20). Pacing started at a cycle length (PCL) of 140 ms. Experiments were run at 1.5× diastolic threshold; voltage was increased if capture failed, unless arrhythmia or ischemia occurred.

### Rapid Pacing Protocol

Pacing cycle lengths were progressively decreased from 140 to 40 ms (steps: 120, 100, 90, 80, 70, 60, 50). Two-second recordings were taken at each step. If arrhythmia occurred, pacing was paused to capture unpaced VT; pacing resumed afterward. Burst pacing was used to terminate episodes lasting >20 minutes.

### Ischemia/Reperfusion Protocol

A 12-minute no-flow ischemia/reperfusion (I/R) protocol was applied. Pacing at PCL100 continued throughout, except during spontaneous VT. Recordings were taken every minute during 12-minute ischemia and reperfusion. Flow was restored post-ischemia and adjusted to maintain pressure ≥50 mmHg. Hearts were weighed and flash frozen post-protocol.

### qPCR and Western Blotting

RNA >200 nt was extracted using the RNeasy Mini Kit (Qiagen, #74104), and cDNA synthesized using the High Capacity cDNA Reverse Transcription Kit (Applied Biosystems, #4368814). qPCR was performed using Fast SYBR Green Master Mix (Applied Biosystems, #4385616), normalized to ribosomal protein L23. Primer sequences are provided below.

Western blots were conducted on left ventricular lysates as described previously (21). Antibodies and dilutions are listed in Table 1. Densitometry was performed using ImageJ. For blots run separately, a shared lysate control was used to normalize across gels. The normalized data were scaled to the ND WT group mean.

**Table 1:** Western Blot antibody details: in order of appearance in paper.

| Antibody | kDa | Species | Dilution | Company (Catalog #) |
| --- | --- | --- | --- | --- |
| AMPKα | 62 | Rabbit | 1:1000 | Cell Signaling (2532) |
| Phospho-AMPKα (Thr172) (40H9) | 62 | Rabbit | 1:1000 | Cell Signaling (2535) |
| Acetyl-CoA Carboxylase | 280 | Rabbit | 1:1000 | Cell Signaling (3662) |
| Phospho-Acetyl-CoA Carboxylase (Ser79) | 280 | Rabbit | 1:1000 | Cell Signaling (3661) |
| Raptor (24C12) | 150 | Rabbit | 1:1000 | Cell Signaling (2280) |
| Phospho-Raptor (Ser792) | 150 | Rabbit | 1:1000 | Cell Signaling (2083) |
| Phospho-DRP1 (Ser637) | 78-82 | Rabbit | 1:1000 | Cell Signaling (4867) |
| Phospho-DRP1 (Ser616) | 78-82 | Rabbit | 1:1000 | Cell Signaling (3455) |
| DRP1 (D6C7) | 78-82 | Rabbit IgG | 1:1000 | Cell Signaling (8570) |
| NaV1.5 (SCN5A) (493-511) | 210 | Rabbit | 1:1000 | Alomone (#ASC-005) |
| Connexin 43 | 39,41,43,44 | Rabbit | 1:1000 | Cell Signaling (3512) |
| ULK1 (D8H5) | 150 | Rabbit IgG | 1:1000 | Cell Signaling (8054) |
| Phospho-ULK1 (Ser555) (D1H4) | 150 | Rabbit | 1:1000 | Cell Signaling (5869) |
| MFN1 | 84, 42 | Rabbit, polyclonal | 1:1000 | Proteintech (13798-1-AP) |
| Mitofusin-2 (D2010) | 80 | Rabbit IgG | 1:1000 | Cell Signaling (9482S) |
| OPA1 (D6U6N) | 78-82 | Rabbit IgG | 1:1000 | Cell Signaling (80471) |
| GAPDH (14C10) | 37 | Rabbit | 1:1000 | Cell Signaling (2118) |

Immunofluorescence
| Antibody | Species | Dilution | Company (Catalog #) |
| --- | --- | --- | --- |
| *Primary antibodies* |  |  |  |
| Anti-N-Cadherin antibody | Mouse | 1:100; 1:50 | Sigma- Aldrich (C3865) |
| Anti-Connexin-43 antibody | Rabbit | 1:2000 | Sigma-Aldrich (C6219) |
| Phospho-Connexin 43 (Ser368) antibody | Rabbit | 1:100 | Cell Signaling (3511) |
| *Secondary* |  |  |  |
| *antibodies* |  |  |  |
| Goat anti-Mouse IgG (H+L) Highly Cross-Adsorbed Secondary Antibody, Alexa Fluor™ 488 | Goat | 1:1000 | ThermoFisher Scientific/Invitrogen (A-11029) |
| Goat anti-Rabbit IgG (H+L) Cross-Adsorbed Secondary Antibody, Alexa Fluor™ 568 | Goat | 1:1000 | ThermoFisher Scientific/Invitrogen (A-11011) |
| Alexa Fluor™ 647 Phalloidin |  | 1:400 | ThermoFisher Scientific/Invitrogen (A22287) Molecular Probes |

### Immunohistochemistry and Immunofluorescence

Fresh hearts were rinsed in PBS, embedded in OCT compound (Fisher #23-730-571), and stored at −80°C. Ventricular cryosections (7 µm) were mounted on coated glass slides (Fisher #12-550-15).

Slides were fixed in 4% paraformaldehyde (Electron Microscopy Sciences #15710-S), washed in PBS, permeabilized with 0.3% Triton-X, and blocked with 5% BSA for 1 hour. Primary antibodies (Table 1) were applied overnight at 4°C. Slides were washed and incubated with secondary antibodies for 1 hour at room temperature, protected from light. Coverslips were mounted with VECTASHIELD HardSet Antifade (Vector #H-1400-10) and stored at 4°C. Confocal imaging was performed using a Leica TCS SP8 inverted microscope at 40× and 63×. Three hearts per group were stained for Cx43 or p-Cx43 (AF568), N-Cadherin (AF488), and F-actin (AF647). No-primary controls were included.

### Colocalization Analysis

Confocal images were analyzed using ImageJ. Thresholding was applied based on no-primary controls. Co-localization was quantified using the JACoP plugin, yielding Pearson’s and Manders’ coefficients.(22) Three images per heart were analyzed.

### Trichrome Staining

Masson’s trichrome staining was performed using the Abcam Trichrome Stain Kit (#ab150686).

### Dihydroethidium (DHE) Staining

Midventricular cryosections (15 µm) were exposed to PBS or 500 µM H2O2 (positive control) for 20 minutes at 37°C. Sections were washed, stained with DHE and Hoechst 33342, and incubated at 37°C for 30 minutes, protected from light (23–25). After washing, slides were mounted with VECTASHIELD and dried. Confocal Z-stacks of the left ventricular free wall were acquired (step size: 0.69 µm, resolution: 2048×2048, 63× objective). Accurate documentation of the timing and sequence of each step in the protocol is essential to ensure comparable results and to eliminate measurement bias in the analysis of the linear increase in DHE-derived fluorescence intensity due to basal ROS activity (25).

### DHE Image Analysis

ImageJ was used for blinded analysis. Corrected total cell fluorescence (CTCF) was calculated as: CTCF = Integrated Density − (Area × Background Intensity). Nuclear signal was isolated using a mean filter, and cytosolic fluorescence was quantified.(26) Background intensity was averaged from five 100×100 pixel areas outside tissue (27). Analysis was automated in MATLAB.

### Transmission Electron Microscopy

TEM of WT and KD hearts (ND-fed) was performed as previously described (18).

### Statistical Analysis

For two-group comparisons, unpaired two-tailed Student’s t-tests were used. Fisher’s exact test was employed to analyze contingency data. One-way ANOVA followed by Tukey’s post hoc test was used for comparisons involving three or more groups. Analyses were performed using GraphPad Prism (v10.11). Data are reported as mean ± standard deviation; p < 0.05 was considered statistically significant.

## Figure Legends

**Supplemental Figure 1:** Prolonged metabolic stress downregulates AMPK activation and loses cardioprotection. **A.** Cumulative weight gain from WT mice on an 8-week (red) vs 16-week (green) high fat diet. **B.** Representative western blots of phospho-AMPK, phospho-Raptor, and GAPDH of 8-week and 16-week HFD mice. **C.** Time course of CV during I/R protocol normalized to basal pre-ischemic CV, showing worsened recovery in the 16-week group (green) compared to the 8-week group (red, also show in Figure 2C and 4B).

## References

1. Holmstrom L, Junttila J, and Chugh SS. Sudden Death in Obesity: Mechanisms and Management. Journal of the American College of Cardiology. 2024;84(23):2308–24.

2. Yao Y, Xue J, and Li B. Obesity and sudden cardiac death: Prevalence, pathogenesis, prevention and intervention. Front Cell Dev Biol. 2022;10:1044923.

3. Prevention CfDCa. Adult Obesity Facts.. Accessed February 12, 2025.

4. Aune D, Schlesinger S, Norat T, and Riboli E. Body mass index, abdominal fatness, and the risk of sudden cardiac death: a systematic review and dose-response meta-analysis of prospective studies. Eur J Epidemiol. 2018;33(8):711–22.

5. Afshin A, Forouzanfar MH, Reitsma MB, Sur P, Estep K, Lee A, et al. Health Effects of Overweight and Obesity in 195 Countries over 25 Years. N Engl J Med. 2017;377(1):13–27.

6. Lavie CJ, Milani RV, and Ventura HO. Obesity and cardiovascular disease: risk factor, paradox, and impact of weight loss. J Am Coll Cardiol. 2009;53(21):1925–32.

7. Carbone S, Canada JM, Billingsley HE, Siddiqui MS, Elagizi A, and Lavie CJ. Obesity paradox in cardiovascular disease: where do we stand? Vasc Health Risk Manag. 2019;15:89–100.

8. Herzig S, and Shaw RJ. AMPK: guardian of metabolism and mitochondrial homeostasis. Nat Rev Mol Cell Biol. 2018;19(2):121–35.

9. Watt MJ, Steinberg GR, Chen ZP, Kemp BE, and Febbraio MA. Fatty acids stimulate AMP-activated protein kinase and enhance fatty acid oxidation in L6 myotubes. J Physiol. 2006;574(Pt 1):139–47.

10. Kim AS, Miller EJ, Wright TM, Li J, Qi D, Atsina K, et al. A small molecule AMPK activator protects the heart against ischemia-reperfusion injury. J Mol Cell Cardiol. 2011;51(1):24–32.

11. Zaha VG, Qi D, Su KN, Palmeri M, Lee HY, Hu X, et al. AMPK is critical for mitochondrial function during reperfusion after myocardial ischemia. J Mol Cell Cardiol. 2016;91:104–13.

12. Russell RR, 3rd, Li J, Coven DL, Pypaert M, Zechner C, Palmeri M, et al. AMP-activated protein kinase mediates ischemic glucose uptake and prevents postischemic cardiac dysfunction, apoptosis, and injury. J Clin Invest. 2004;114(4):495–503.

13. Yashirogi S, Nagao T, Nishida Y, Takahashi Y, Qaqorh T, Yazawa I, et al. AMPK regulates cell shape of cardiomyocytes by modulating turnover of microtubules through CLIP-170. EMBO Rep. 2021;22(1):e50949.

14. Saffitz JE, Laing JG, and Yamada KA. Connexin expression and turnover: implications for cardiac excitability. Circ Res. 2000;86(7):723–8.

15. Hesketh GG, Shah MH, Halperin VL, Cooke CA, Akar FG, Yen TE, et al. Ultrastructure and regulation of lateralized connexin43 in the failing heart. Circ Res. 2010;106(6):1153–63.

16. Akar FG, Spragg DD, Tunin RS, Kass DA, and Tomaselli GF. Mechanisms underlying conduction slowing and arrhythmogenesis in nonischemic dilated cardiomyopathy. Circ Res. 2004;95(7):717–25.

17. Tocchetti CG, Caceres V, Stanley BA, Xie C, Shi S, Watson WH, et al. GSH or palmitate preserves mitochondrial energetic/redox balance, preventing mechanical dysfunction in metabolically challenged myocytes/hearts from type 2 diabetic mice. Diabetes. 2012;61(12):3094–105.

18. Cacheux M, Rudokas M, Tieu A, Rizk JA, Hummel ME, and Akar FG. Quantitative Assessment of Mitochondrial Morphology and Electrophysiological Function in the Diabetic Heart. Methods Mol Biol. 2024;2803:75–86.

19. Cacheux M, Strauss B, Raad N, Ilkan Z, Hu J, Benard L, et al. Cardiomyocyte-Specific STIM1 (Stromal Interaction Molecule 1) Depletion in the Adult Heart Promotes the Development of Arrhythmogenic Discordant Alternans. Circ Arrhythm Electrophysiol. 2019;12(11):e007382.

20. Raad N, Bittihn P, Cacheux M, Jeong D, Ilkan Z, Ceholski D, et al. Arrhythmia Mechanism and Dynamics in a Humanized Mouse Model of Inherited Cardiomyopathy Caused by Phospholamban R14del Mutation. Circulation. 2021;144(6):441–54.

21. Toksoy Z, Ma Y, Goedeke L, Li W, Xiaoyue H, Wu X, et al. Role of AMPK in Atrial Metabolic Homeostasis and Substrate Preference. 2024.

22. Bolte S, and Cordelières FP. A guided tour into subcellular colocalization analysis in light microscopy. J Microsc. 2006;224(Pt 3):213–32.

23. Mesubi OO, Rokita AG, Abrol N, Wu Y, Chen B, Wang Q, et al. Oxidized CaMKII and O-GlcNAcylation cause increased atrial fibrillation in diabetic mice by distinct mechanisms. J Clin Invest. 2021;131(2).

24. Wang Q, and Zou MH. Measurement of Reactive Oxygen Species (ROS) and Mitochondrial ROS in AMPK Knockout Mice Blood Vessels. Methods Mol Biol. 2018;1732:507–17.

25. Sözer EB, Semenov I, and Vernier PT. Dihydroethidium-derived fluorescence in electrically stressed cells indicates intracellular microenvironment modifications independent of ROS. Bioelectrochemistry. 2024;160:108751.

26. Kelley JB, and Paschal BM. Fluorescence-based quantification of nucleocytoplasmic transport. Methods. 2019;157:106–14.

27. Ansari N, Müller S, Stelzer EHK, and Pampaloni F. In: Conn PM ed. Methods in Cell Biology. Academic Press; 2013:295–309.

