## Supplementary figures and images for "Transient AMPK activation by nutrient stress of high fat diet preserves cardiac electrophysiological stability and protects against arrhythmias"

### SUPPLEMENTAL FIG 1

#### Supplemental Figure 1

A.

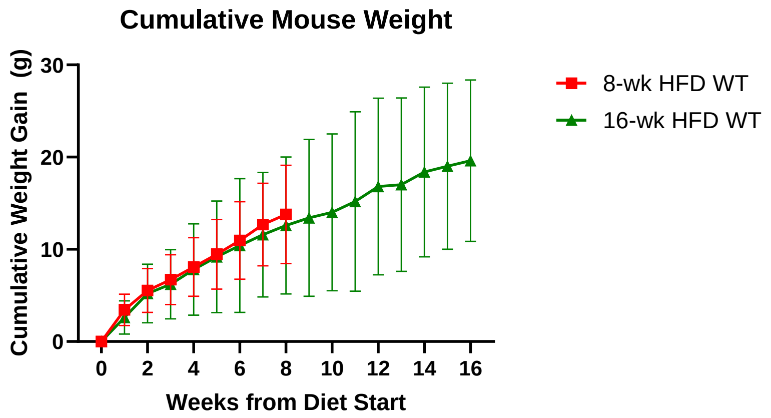


B.

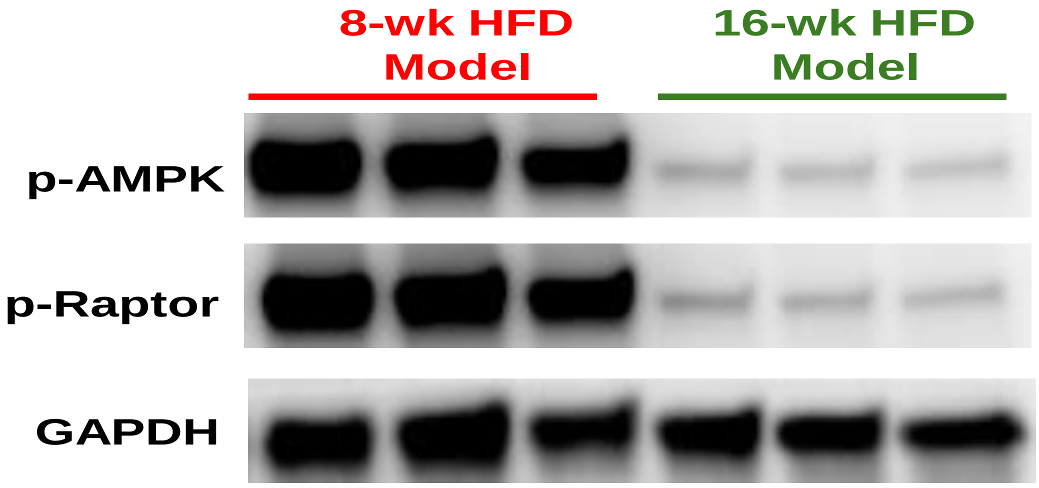


C.

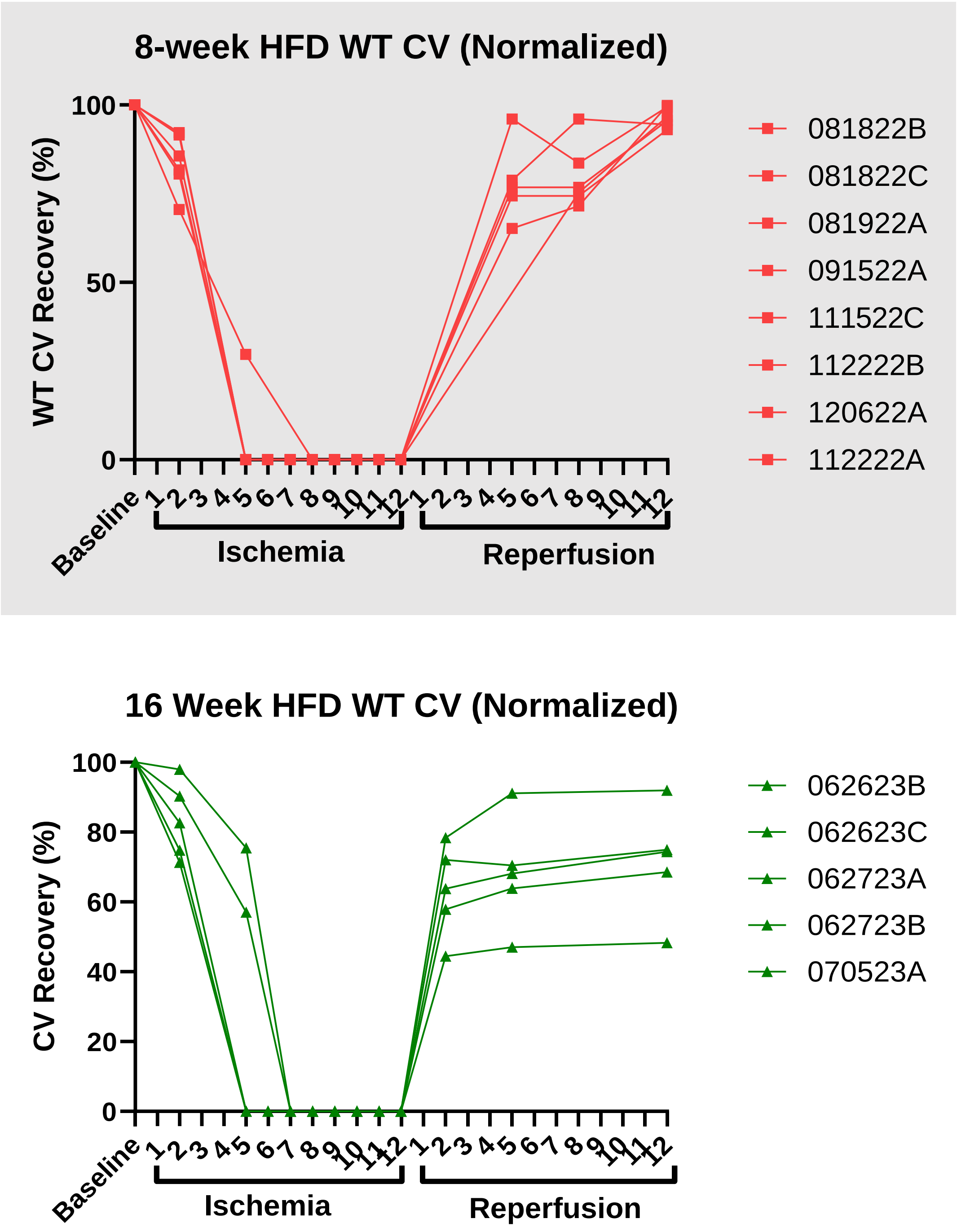
